# Genome-wide signatures of reproductive isolation shape the varied genomic landscape of the roundtail horned lizard (*Phrynosoma modestum*)

**DOI:** 10.1101/2025.09.25.678555

**Authors:** Julia Amoroso, Keaka Farleigh, Erika Crispo, Christopher Blair

**Affiliations:** Biology PhD Program, CUNY Graduate Center, 365 5th Ave., New York, NY 10016, USA; University of Virginia, Charlottesville, VA 22904, USA; Department of Biology, Pace University, One Pace Plaza, New York, NY 10038, USA; Department of Biological Sciences, New York City College of Technology, The City University of New York, 285 Jay Street, Brooklyn, NY 11201, USA

**Keywords:** *Phrynosoma modestum*, islands of divergence, allopatry, speciation

## Abstract

Population divergence is promoted and inhibited by gene flow and divergent selection, but the mechanisms of and relationship between these two processes remain poorly understood. Modelling the selective pressures at play in a natural population requires a thorough understanding of species structure and demographic history, which can be helpful in hypothesizing the genetic and evolutionary factors that underlie divergence. In this study, we assess whole genome sequences of round-tailed horned lizards (*Phyrnosoma modestum)* from throughout the species range and combine phylogenetic analyses with genetic landscape scans to understand how current genetic diversity has been influenced by demographic histories and evolutionary pressures. Maximum likelihood phylogenetic analysis supports two lineages within the species, correlating with a North/South population divide that likely diverged about 7 Ma and displayed little migration. Intermediate *gdi* values indicate that the two lineages may be in the gray area of speciation, yet our results also support significant isolation-by-distance (IBD). Negative values of Tajima’s *D* offer support for selection acting on *P. modestum*, but may stem from recent population expansions. Genomic diversity within populations and islands of divergence between populations are also detected across the *P. modestum* genome, specifically exhibiting differentiation patterns linked to reproductive isolation and within-population selection. We posit potential explanations for the genomic landscape characterized here, namely that allopatry plays a large role in shaping genomic divergence across the species’ range. Taken together, our results provide a picture of a species currently maintaining species integrity despite significant genomic signatures of geographic structure and reproductive isolation.

## Introduction

Intraspecific variation is required for evolution through natural selection. In natural populations, intraspecific variation is generated and influenced by evolutionary processes closely linked with population divergence. Genetic drift and divergent selection increase divergence between populations and, over time, result in the evolution of reproductive isolation (Kimura, 1968; King & Jukes, 196; Nei et al., 1983). Contrastingly, gene flow and balancing selection decrease divergence and inhibit the evolution of reproductive isolation (Wu, 2001). Yet, our understanding of these processes, the genomic regions they influence, and the genes they involve remains incomplete, even as it is a central goal of evolutionary biology.

Key to our current understanding of how population divergence develops has been hypothesizing models of divergence, which can produce recognizable genetic signatures and lead to genomic islands of differentiation (Shang et al. 2023). These islands are genomic regions that display high levels of relative differentiation (Fst) in comparison to the background rate, therefore assumed to reflect loci under or linked to those under divergent selection (Han et al., 2017; Wu, 2001). Pairing Fst with measurements of absolute sequence divergence between populations (dxy) and nucleotide diversity within populations (π) allows for further predictions about the presence of gene flow and the type of selection acting on regions of divergence (Cruickshank & Hahn, 2014; Turner et al., 2005). Irwin et al. (2018) propose four models to explain common patterns of population statistics within genomic regions of divergence (Figure 1). In a divergence with gene flow model (red), two populations are in physical contact (sympatric or parapatric) and gene flow occurs between them. Genomic regions that contribute to reproductive isolation act as barriers to gene flow, exhibiting increased Fst and dxy and decreased π due to selection against gene flow. Gene flow can still occur in neutral regions, leading to lower Fst and dxy in these portions of the genome, and increased π as selection is not acting against gene flow. The selection in allopatry model (blue) identifies increased regions of Fst due to within-population selection. Within-population positive or background selection reduces π in genome regions experiencing selection, increasing Fst in these regions; balancing selection is expected to increase π. Dxy remains similar between regions under selection and neutral regions because it is unaffected by π (Cruickshank & Hahn, 2014). A recurrent selection model (yellow) is similar but also invokes selection in the ancestral population, which decreases dxy and π even more than the selection in allopatry model since selection occurs in both the ancestral and modern populations. In a geographic sweep before selective differentiation model (purple), advantageous alleles have spread throughout a geographically structured species complex (reducing dxy in those genomic regions). Geographic stratification then promotes selection within populations, decreasing π and increasing Fst if different advantageous alleles are selected for in different populations (Irwin et al., 2018).

**Figure 1.**
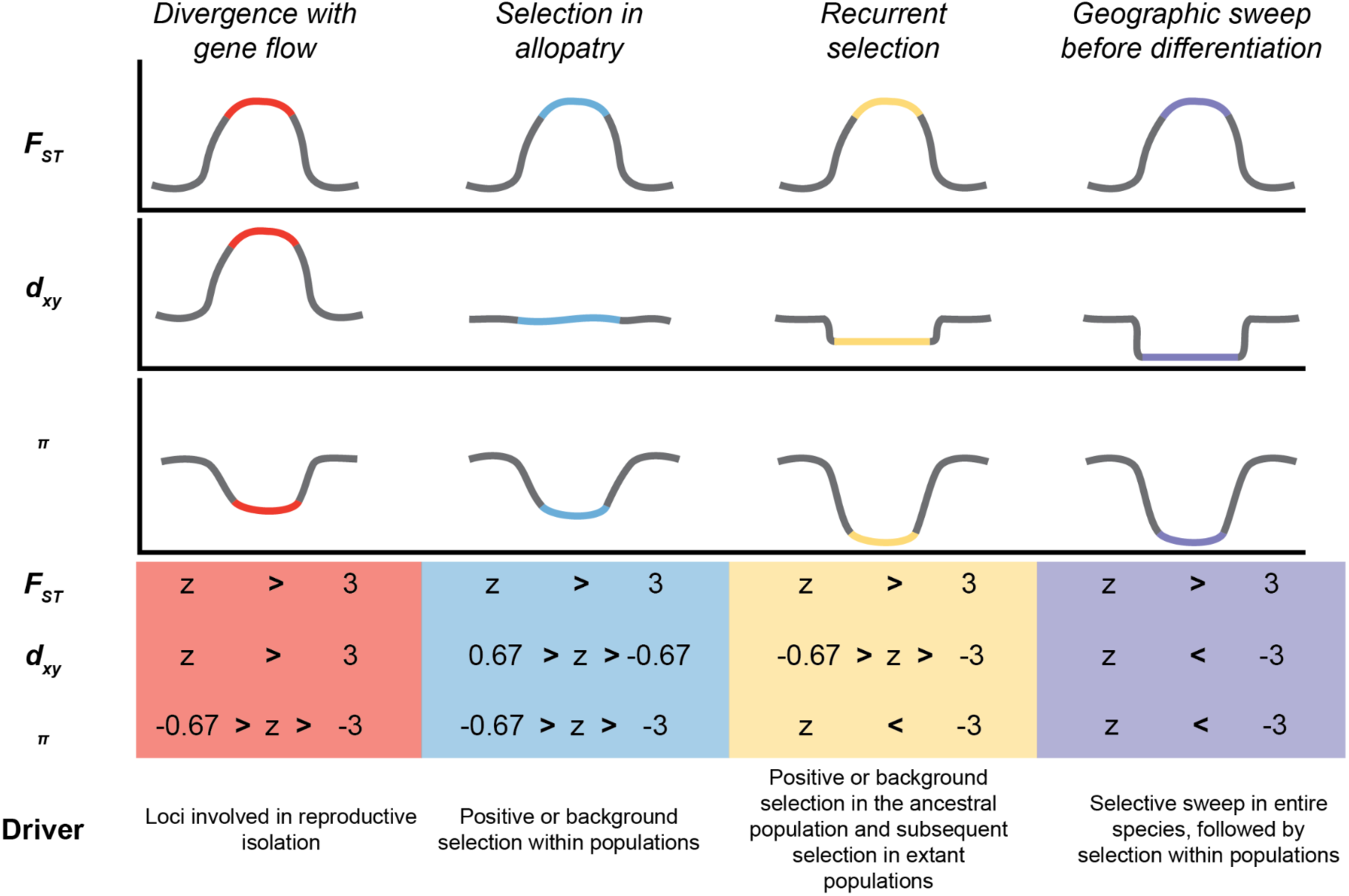
Conceptual visualization of different scenarios that form genomic divergence islands. The illustrations show the relationship between the genomic divergence island (colored region) and the genomic background (gray). The thresholds we used to identify candidate windows are in the colored boxes and the source of selection in each scenario is listed at the bottom. These scenarios and the sources of their selection were redrawn from Irwin et al. (2018).

Correctly analyzing the signal of islands of divergence can be clouded by the effects of genetic architecture and long-term evolutionary dynamics (Nosil et al., 2009). Reduced recombination can play an outsized role in shaping a heterogeneous genetic landscape, but understanding historic gene flow between populations helps distinguish whether genetic differentiation has also been formed by selection (along with incorporating absolute metrics of divergence in island identification, see Noor & Bennett, 2009; Han et al., 2017; Martin et al., 2013; Renaut et al., 2013). Historical introgression events between ancestral lineages can also be difficult to distinguish from present ongoing gene flow (Pavón-Vázquez et al., 2024).

Linking models of selective pressures in islands of divergence with a phylogeographic examination of species history and structure deepens our understanding of how and why selective pressures develop in natural populations (Brown et al., 2016; Blair et al., 2026; Edwards et al., 2021; Ravinet et al., 2017). Phrynosomatid lizards serve as an ideal system to connect models of genomic divergence with broader-scale evolutionary histories, as recent genomic work in the *Phrynosoma* (horned lizard) genus has largely resolved phylogenetic relationships between members of the group (Leache et al., 2021). Within *Phrynosoma,* species-level evolutionary histories and population structure have been examined for many species, yet round-tailed horned lizards have been notably excluded (Farleigh et al., 2021; Finger et al., 2021; Gottscho et al., 2023). The round-tailed horned lizard (*Phrynosoma modestum*) is a member of the *Doliosaurus* clade of *Phrynosoma* and diverged from its sister species *Phrynosoma platyrhinos* and *Phrynosoma goodei* roughly 15 million years ago (Ma; Leaché & Linkem, 2015; Leaché & McGuire, 2006). Recent ecological and genomic work on *P. modestum* has been sparse, likely due to its small size and cryptic coloration (Whiting & Dixon, 1996). Species distribution estimates for *P. modestum* range across the mesa region of southeastern Arizona into western and southern Texas, extending down to northern Mexico (Weese, 1917; Whiting & Dixon, 1996). Numerous geographic features divide this range, including the Rocky Mountains, Rio Grande, and other small water features (Bundy & Neess, 1958). Little is understood about species behavior and evolutionary history. However, in an experimental behavioral study, members of the species displayed a smaller home range than expected under random conditions, and have presented phenotypic variation in dorsal color (Munger, 1984b). While this presents compelling evidence for the possibility of population differentiation across the species range, further investigation should verify if and how genetic divergence underlies *P. modestum*’s phenotypic variation.

In this study, we assess genetic variation and divergence across the native range of *P. modestum* to explore the selective pressures and demographic histories shaping current population diversity. The observation of phenotypic variation, coupled with significant geographic barriers across the species range and evidence for limited home range sizes, led us to hypothesize that the *P. modestum* genome will display genomic divergence due to selective pressures linked to reproductive isolation and within-population selection. We analyze multilocus and SNP datasets based on whole-genome sequences of *P. modestum*. We carry out population structure analyses and reconstruct demographic histories using multiple tree-based methods. To assess selection pressures causing genetic divergence within *P. modestum*, we test for signatures of each of four models of islands of divergence and perform gene ontology analyses for genes in proximity of diverging genomic windows.

## Methods

### Data collection

We obtained data from 26 *Phrynosoma modestum* individuals and one *P. douglasii* individual with geographic coordinate data (Table S1). Tissue samples were obtained from museums and field sampling of the Southwestern United States and Mexico. All field work was conducted under IACUC approval from Brooklyn College (Protocol # 312). The *P. douglasii* individual was included as an outgroup. We extracted DNA from samples using a Zymo Quick-DNA Miniprep Kit (Zymo Research) following manufacturer’s protocols. DNA quantity was measured using a Qubit 4.0 fluorometer (Invitrogen) following manufacturer’s recommendations. DNA quality was assessed by running 3 ul of DNA extract on a 1% agarose gel stained with SYBR Green (Thermo Fisher Scientific). Samples that passed QC were sent to Novogene for whole genome library preparation and sequencing.

### Data sequencing, assembly, and filtering

Novogene generated whole genome data on an Illumina NovaSeq6000 using 150 bp paired-end sequencing for 27 individuals (26 *P. modestum*, 1 *P. douglasii*) and performed initial bioinformatic analyses. Sequence data were mapped to the publicly available reference genome of *P. platyrhinos* (Koochekian et al., 2022) using the Burrows-Wheeler Aligner (BWA; Li et al., 2009a) and SNPs were called and filtered using SAMtools (Li et al., 2009b) and GATK v4.6.1.0 (DePristo et al., 2011). We applied additional filters for population genetic analyses with vcftools v0.1.16 (Danecek et al., 2011), requiring at least 90% data, a minimum quality score of 20, SNPs that were biallelic, and a minor allele frequency of 0.05. SNPs were thinned by 10kb to minimize linkage disequilibrium. One ingroup individual was removed due to high levels of missing data (see Table S2).

We performed additional mapping, SNP calling, and filtering for sliding-window analyses since dxy and π calculations require both invariant and variant sites (Korunes & Samuk, 2021). We followed a similar pipeline to Novogene of mapping with BWA, but performed SNP calling in bcftools v1.21 (Danecek et al., 2021) and did not apply a minor allele frequency or minimum allele filters. Finally, we used the captus v1.1.0 pipeline (Ortiz et al., 2023) to assemble sequence reads into loci. We followed the captus tutorial (https://edgardomortiz.github.io/captus.docs/) and extracted tetrapod ultra-conserved elements (UCEs; 5,472 baits for 5,060 loci; Tetrapods-UCE-5Kv1 from https://www.ultraconserved.org/; Faircloth et al., 2012) to represent our loci. We used default settings for each step, except for assembly, where we required that contigs had a minimum depth of 5. We chose the bait with the least missing data if multiple baits were identified for a single locus. We also used captus to extract the mitochondrial genome (mitogenome), using the *P. blainvilli* mitogenome (NCBI RefSeq: NC_036492.1) as the reference (Ayala et al., 2017).

### Population structure and demographic history

We inferred the number of genetic clusters (*K*) and ancestry of each *P. modestum* sample using ADMIXTURE v1.3.0 (Alexander et al., 2009). ADMIXTURE input files were generated from our filtered ingroup VCF using PLINK v1.9.0b.7.7 (Purcell et al., 2007), which provided us with the input bed file. We determined the best *K* for our dataset by calculating the cross-validation error for *K* values ranging from 1 to 8 and selecting the *K* with the lowest error (Figure S1). We generated ancestry plots using our optimal *K* in PopGenHelpR v1.3.2 (Farleigh et al., 2025).

We used IQ-TREE v2.2.2.6 (Minh et al., 2020) to run a partitioned maximum likelihood (ML) phylogenetic analysis on the tetrapod loci from the captus assembly, rooted using *P. douglasii*. ModelFinder (Kalyaanamoorthy et al., 2017) was used to find the best substitution model using the Bayesian information criterion (BIC) for each partition and to merge partitions (MFP+MERGE command) if it improved the overall model fit; initially, each UCE locus was considered a partition. ModelFinder uses a greedy algorithm that first determines the best fit substitution model for individual partitions and then tests whether or not merging two partitions and assigning them the same substitution model improves the overall model fit (Lanfear et al., 2012; Minh et al., 2020). We evaluated branch support using ultrafast bootstrapping implemented by UFBoot2 (Hoang et al., 2018) and non-parametric Shimoda-Hasegawa approximate likelihood ratio tests (SH-aLRTs) using 10,000 ultrafast bootstraps and 1,000 SH-aLRTs. We also ran a partitioned ML analysis of mitogenomes with IQ-TREE v2.2.2.6 using the same parameters as the UCE dataset (ModelFinder, 10,000 UFboostraps; 1,000 SH-aLRTS) and defining initial partitions by gene.

Finally, we ran a fixed species delimitation and species tree analysis (A00) in BPP v4.8.4 (Flouri et al., 2018) to estimate the demographic history of *P. modestum*. We used the recently developed MSC-M model to estimate population sizes (θ) for both ancestral and current populations, along with divergence times (τ) and effective migration rates (ɷ) between populations (Flouri et al. 2023). Note that BPP estimates mutation-scaled migration rates with ɷ = m/μ. Analyses were performed with inverse gamma priors for θ (3, 0.004) and τ (3, 0.04), and a gamma prior for ɷ (2, 1). We specified two migrations bands, one from lineage A to lineage B and vice versa. We used a total of 4,961 UCE loci from all samples in each population and one outgroup sample, and performed four independent runs to ensure convergence and adequate mixing. Runs used a burn-in of 50,000, a sample frequency of 2, and 100,000 total samples. Tracer v1.7.2 (Rambaut et al., 2018) was used to assess convergence, mixing, and ESS values (target >200) and also to combine results across runs. To convert ɷ to number of migrants per generation (M), we used the following equation:

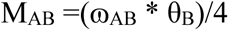

BPP estimates divergence times in units of expected substitutions per site. To convert τs to absolute time in millions of years, we first assumed a divergence time of 15 Ma between *P. modestum* and *P. douglasii* based on the results of a previous phylogenomic study (Leache & Linkem, 2015). We then used the formula μ = τ/T to calculate a substitution rate for our UCE loci. Using the calculated μ, we rescaled the remaining τs to millions of years.

To provide further support for the independence of lineages and account for potential gene flow, we calculated the genealogical divergence index (*gdi*; Jackson et al. 2017) following Pavon-Vazquez et al. (2024). Specifically, we used BPP to simulate 1 million gene trees with two species (A and B) and three alleles (x_1_, x_2_, y) using demographic parameters estimated from our combined runs of the empirical data. Importantly, this calculation explicitly accommodated migration. We calculated *gdi* as the proportion of simulated gene trees with topology G_x_ = ((x_1_, x_2_), y) where the coalescence time of x_1_ and x_2_ was more recent than the population divergence time τ (Kornai et al. 2024). Two sets of simulations were performed to calculate *gdi* for both lineages. We followed recent guidelines for interpretation of *gdi* values, with estimates <= 0.3 indicating conspecificity, values >= 0.7 suggesting different species, and intermediate values inconclusive (Jackson et al. 2017; Chambers et al. 2025).

### Intraspecific variation

We estimated the genetic diversity and differentiation in our data using PopGenHelpR v1.3.2. The observed heterozygosity of each population and the proportion of heterozygous loci in each individual was estimated using the *Heterozygosity* function. The genetic differentiation between populations (Fst) was calculated using the *Differentiation* function.

We also evaluated the correlation between geographic and genetic distance to test for evidence of isolation-by-distance (IBD) in our dataset using the R packages nlme v3.1-165 (Pinheiro et al., 2012) and corMLPE v0.0.3 (Pope, 2021). These packages use Clarke’s maximum likelihood population effects model to identify a correlation between genetic and geographic distance (Clarke et al., 2002).

### Population divergence

We generated sliding-window summary statistics of genomic diversity and differentiation in pixy v1.2.11.beta1 (Korunes & Samuk, 2021) to understand how selection may be promoting genetic divergence between populations following the conceptual models of Irwin et al. (2018). For every non-overlapping 10 kb window across the *P. modestum* genome, we calculated nucleotide diversity within both populations (π), along with absolute nucleotide divergence (dxy) and relative genetic differentiation between populations (Fst).

To make comparisons between genomic windows, we standardized per-window diversity metrics by calculating z-scores for each metric. We identified Fst peaks as windows with a z-score of three or above (Stone & Wessinger, 2024). We identified regions with a lower π and dxy than the genomic background as regions with a z score between −3 and −0.67 (see divergence with gene flow, selection in allopatry, recurrent selection in Fig. 1).We evaluated patterns of π in two ways. First, we calculated measures of π within each population instead of averaging across populations to identify genomic windows exhibiting patterns of different selective models in each population. Second, we averaged π across populations to calculate z-scores and identify outlier windows across populations. We identified regions with an average dxy as windows with a z-score between −0.67 and 0.67 (see selection in allopatry, Fig. 1). Both invariant and variant sites were used for this analysis (no minor allele frequency filter) since π and dxy calculations require both types of sites (Korunes & Samuk, 2021). We also calculated Tajima’s *D* for each 10kb window using vcftools v0.1.15 to test if the proposed windows exhibited differences in allele frequencies. We compared Tajima’s *D* values across different selection models in each population using Kruskal-Wallis and pairwise Wilcoxon tests to determine if they were significantly different in R (R Core Team, 2023).

### Gene ontology

We extracted genes within highly differentiated windows of the *P. modestum* genome to understand which genes may be linked to windows of divergence. We used bedtools v2.31.10 (Quinlan & Hall, 2010) to intersect genomic windows of divergence in *P. modestum* with the annotated *P. platyrhinos* genome (Koochekian et al., 2022) and isolated known genes within each window. We extracted fasta files of known genes using bedtools and performed a nucleotide BLAST with *Sceloporus undulatus* as the reference database to identify common gene names (Altschul et al., 1990). *Sceloporus undulatus* was chosen for its well-notated BLAST entries. STRING v12.0 (Szklarczyk et al., 2025) was then used to understand potential functions and associations between known genes in windows of divergence.

## Results

### SNP filtering

After sequencing, we obtained a mean of 149,094,524 reads per sample. An average of 139,569,010 reads mapped to the *P. platyrhinos* reference genome, representing 93.57% of the genome (Table S2). After initial variant calling and filtering performed by Novogene, our full dataset contained 45,245,582 SNPs. Additional filtering to generate the final ingroup VCF produced 15,368 SNPs from 25 *P. modestum* individuals for population genetic analysis. The sliding-window dataset of variant and invariant sites contained 1,540,051,645 sites. After captus assembly, we recovered an average of 4,986 tetrapod UCE loci per sample with an average depth of 8.4 (Table S3).

### Population structure and history

Cross-validation error in ADMIXTURE was minimized at *K* = 2, yielding two major genetic clusters within *P. modestum*. The clusters corresponded to a North population (lineage A) spanning New Mexico and Texas and a South (lineage B) population in Mexico (Fig. 2A). All samples at the geographic extremes of the North and South populations displayed low levels of admixture. Four individuals in Southern Texas displayed mixed ancestry, where the proportion of ancestry from each population increased with proximity to either cluster (Fig. 2A,D).

**Figure 2.**
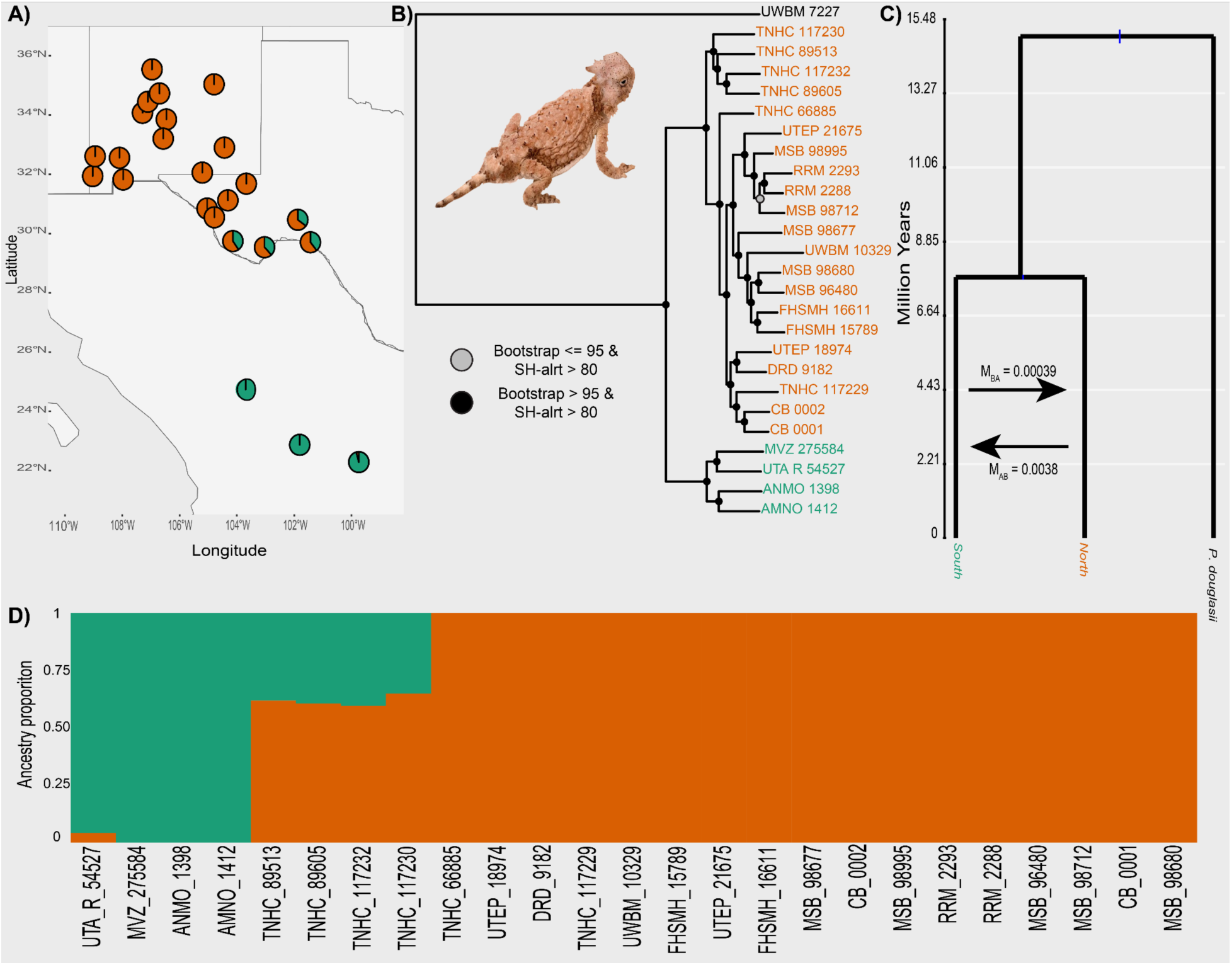
A & D) ADMIXTURE results clustering samples into two populations. Each pie chart and vertical bar represents an individual (in A & D, respectively) with the proportion of each color indicating the proportion of ancestry from a particular cluster. B) A maximum-likelihood genealogy inferred using a partitioned analysis in IQ-TREE. Node values are represented with colored dots indicating the ultrafast bootstrap support and Shimoda-Hasegawa approximate Likelihood Ratio Tests (SH-aLRTs). Black dots indicate ultrafast bootstrap support over 95, while gray indicates less than or equal to 95. Both colors indicate SH-aLRTs values above 80. C) Divergence time and migration rate estimates from BPP MSC-M analysis.

Our ML phylogenetic analysis of the UCE loci strongly supported two lineages corresponding to the ADMIXTURE populations (Fig. 2B). The four Southern Texas admixed individuals formed a divergent cluster but were ultimately placed within the North population lineage, concordant with their ancestry make-up. ModelFinder identified 189 partitions, with an average of 761 parsimony informative sites and 7,910,704 total sites. The ML phylogenetic analysis of the mitogenome sequences was concordant with the UCE loci, supporting two lineages corresponding to the ADMIXTURE populations. The mitogenome ML phylogenetic analysis identified 3 partitions with an average of 479 parsimony informative sites and 15,485 total sites.

Convergence and mixing of the four BPP runs was adequate based on trace plots and ESS values. Contemporary population sizes for the two lineages were similar and larger than the ancestral size (Table S4). The two lineages diverged approximately 7 Ma (Fig. 2C). The calculated substitution rate for the UCE loci was 0.0004 substitutions per site per million years, approximately half the rate as for four-fold degenerate sites and estimates from GBS data (Perry et al., 2018, Pavon-Vazquez et al., 2024). This lends support to the hypothesis that UCE loci may be under some sort of purifying selection that reduces the neutral mutation rate of linked sites. The number of migrants per generation was << 1 for each comparison. However, estimated values were 10x higher from lineage A (North) to lineage B (South) than in the opposite direction (Fig. 2C). The calculated *gdi* for lineage A was 0.397 and the *gdi* for lineage B was 0.443, both falling within the broad zone of uncertainty.

### Intraspecific variation

We found that the North population had an average observed heterozygosity of 0.28 and the South had a heterozygosity of 0.44. The mean proportion of heterozygous sites per individual was 0.31 (Table S6). Fst between the two populations was 0.08. Additionally, a statistically significant correlation between geographic and genetic distance was found, supporting a pattern of IBD in our dataset (p-value < 0.001; rho 0.43).

### Genetic divergence

We used relative and absolute metrics of genetic diversity within populations and differentiation between populations to identify non-overlapping 10 kb windows of divergence and their potential drivers. In the North population, we found 34 candidate windows containing 13 characterized genes associated with a divergence with gene flow model and 85 candidate windows containing 33 characterized genes associated with a selection in allopatry model (Table S5). In the South population, we found 27 candidate windows containing 11 known genes associated with a divergence with gene flow model and 90 candidate windows containing 30 known genes associated with a selection in allopatry model (Table S5). 12 candidate windows containing six known genes were linked to a divergence with gene flow model in both populations. No candidate windows were found with characteristics of recurrent selection or geographic sweep before differentiation (Scenarios 3 and 4). Averaging π identified 34 genomic windows linked to a divergence with gene flow model, and 85 genomic windows associated with a selection in allopatry model, all of which were identified in comparisons without averaging π; therefore, we focused on the within population π windows.

We also tested for evidence of selection in these windows using Tajima’s *D*. Negative values of Tajima’s *D* indicated an excess of rare alleles, which could be linked to a selective sweep or population expansion after a bottleneck. Positive values indicate a lack of rare alleles that could be due to balancing selection or population decrease. A value of 0 indicates that the genomic variation is as expected under a mutation-drift equilibrium. The mean Tajima’s *D* for divergence with gene flow and selection in allopatry windows for the North population was −1.34 and −1.33, respectively (Fig. 3). The mean Tajima’s *D* for divergence with gene flow and selection in allopatry windows for the South population was −0.11 and −0.21, respectively (Fig. 3). The mean Tajima’s *D* for the genomic background (non-candidate windows) was −0.71 and - 0.08 for the North and South populations, respectively (Fig. 3). Kruskal-Wallis tests identified statistically significant differences between different window types for both populations (North: p-value < 2.2e-16; South: p-value = 0.0002). Post-hoc pairwise Wilcoxon tests identified significant differences between divergence with gene flow and neutral windows (p-value = 2.4e-15) and selection in allopatry and neutral windows (p-value < 2e-16) in the North population. The only significant comparison in the South population was between selection in allopatry windows and neutral windows (p-value = 0.0002).

**Figure 3.**
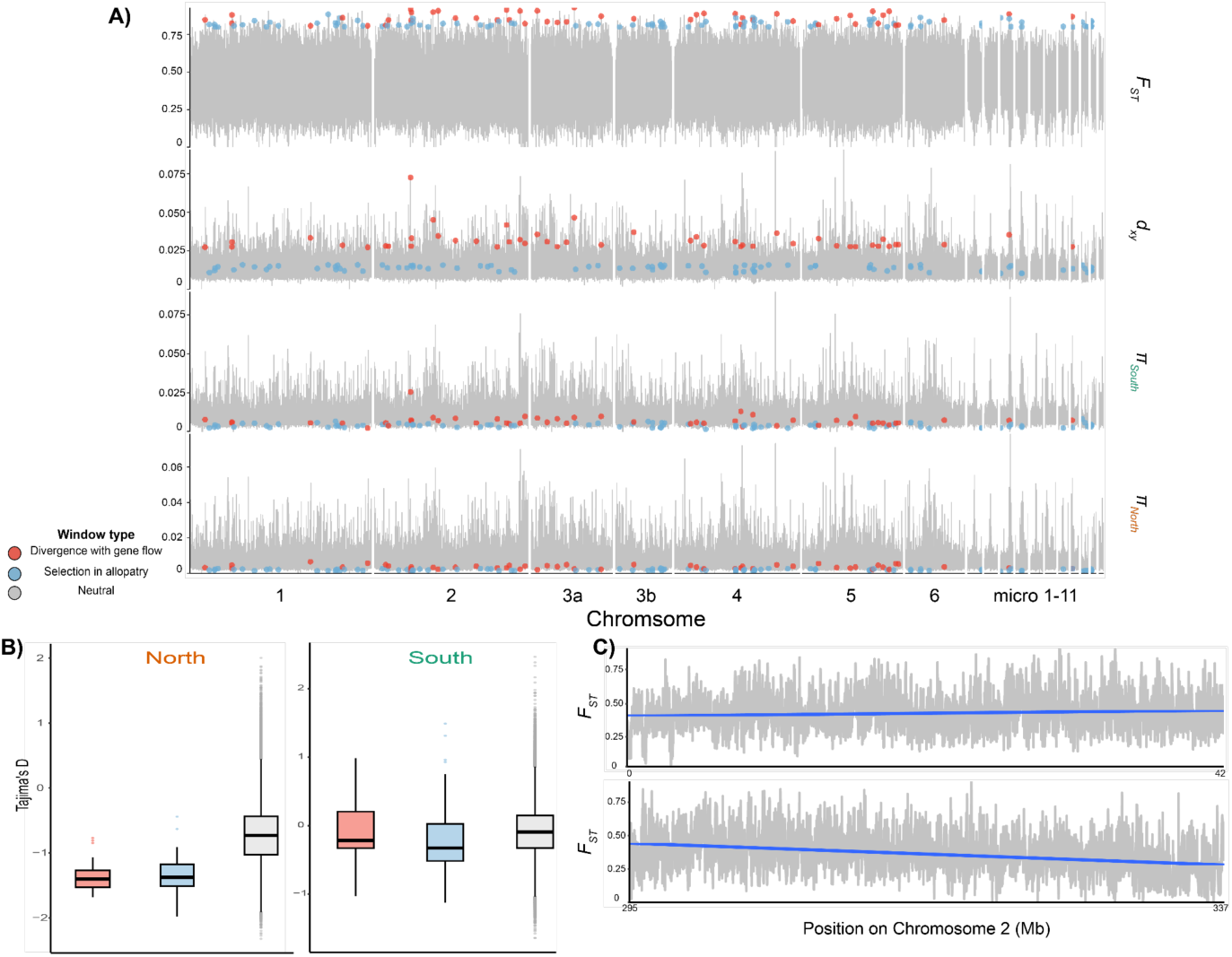
A) Sliding-window divergence and diversity statistics from pixy visualizing genomic divergence islands in divergence with gene flow (red dots) and selection in allopatry (blue dots) scenarios (following Figure 1) across the genome. Genomic windows that did not follow any of the Figure 1 scenarios are labelled as neutral and colored gray; the statistic for each row is indicated on the right. Note that for nucleotide diversity (lower two rows), windows are colored in both populations even though they may only be considered a candidate window in one population (e.g., Chromosome 2 in the south population). B) Tajima’s *D* estimates for the different scenarios observed in our data versus the “neutral” genomic background that did not follow any of the patterns in Figure 1. C) Fst on the first (top) and last (bottom) 42 megabases of Chromosome 2, values are rounded to the nearest whole number. The blue line is a best-fit line from a linear model, which was significant for both ends of the chromosome (beginning p-value = 1.25e-10; end p-value < 2.2e-16), indicating lower Fst on chromosome ends.

Gene ontology analyses of genes extracted from divergent windows identified characterized functions of 51 total genes and functional interactions between seven genes (Table S5). For windows in both populations showing signatures of selection in allopatry, a functional interaction was identified concentrated around three F-Box genes, FBXL20 on Chromosome 10, and FBXO11 and MSH6 on Chromosome 1, along with an interaction between KANSL1 on Chromosome 10 and BRD8 on Chromosome 3. Windows of divergence with gene flow in the South population displayed a functional interaction between DHX29 on Chromosome 3 and NOP14 on Chromosome 8.

## Discussion

Although gene flow is increasingly recognized as a factor in generating intraspecific genetic diversity, allopatric barriers and geographic distance maintain an important role as obstacles to gene flow and promoters of population divergence, particularly for species that display restricted home ranges (Feder et al., 2012; Emelianov et al., 2004; Dool et al., 2021; Munger 1984b, Blair et al., 2026). Population structure and demographic analyses for *P. modestum* suggest a species that, despite maintaining intraspecific genetic diversity through low gene flow, is composed of two highly divergent lineages in the face of geographic distance. Genomic and mtDNA lineage trees are concordant, suggesting the species has not undergone recent hybridization or introgression events with closely related and geographically concurrent species such as *P. goodei* (Jezkova et al., 2011; Leaché & Linkem, 2015).

Demographic analyses estimate that the two lineages, a North and South population, diverged in the Late Miocene, roughly 7 Ma, and display low rates of migration based on UCE loci. Although this is a relatively old divergence time between lineages, *gdi* estimates for both lineages fell into a broadly inconclusive zone and we therefore cannot recommend a full taxonomic distinction between the two lineages. This uncertainty may be due to a physical gap in our sampling in the region of northern Mexico (Figure 2A), and greater sampling efforts at the contact zone between the two populations will help determine potential species boundaries between the lineages (Chambers et al. 2025). The rate of migration from the North population to the South population is ten fold greater than the South-North migration rate, suggesting that Northern alleles are more readily incorporated into the South population and that the populations could have relied on different strategies to persist in their local environments. Phenotypic plasticity, for example, could make it easier to colonize a warmer microhabitat than a colder one (e.g., South population herein; Jezkova et al., 2016). Contrastingly, if the North population has persisted through local adaptation, natural selection would act against any maladapted combinations of alleles to reduce variation in the population and gene flow from South to North. Indeed, although a slightly larger population size is suggested in the North, we observe a lower heterozygosity with an average of 0.28 (Table S6), and certain combinations of alleles may have been promoted by natural selection in that environment. However, this could also be a signature of range expansion or genetic drift. Heterozygosity is higher in the South population, which is unsurprising as we detect more migration from the North into the South. Increased variation may be due to the South population having closer geographic proximity to the ancestral source of the range expansion or a decreased reliance on local adaptation to persist, which would lessen selection against potentially maladapted combinations of alleles. However, the extremely low overall migration rates between these two concurrent and related populations also point to the traditional role assumed by allopatry in promoting population divergence within *P. modestum*.

Overall genomic heterogeneity across the *P. modestum* genome is largely shaped by high signals of divergence at a small number of genomic regions, with 245 10 kb windows following expectations of any model of divergence. Of the four models of selection pressures shaping windows of divergence, outlier windows only display characteristics linked to selection in allopatry and divergence with gene flow (Figure 3A). This supports our hypothesis that, despite geographic structure, genomic divergence will be relatively limited because samples represent two populations within the same species (Irwin et al., 2016). Between the two models of islands of divergence, candidate windows largely support a selection in allopatry model (33 genes in total). While relatively fewer than the previous model, some windows of divergence support a divergence with gene flow model (18 genes in total).

Certain genomic regions, such as the ends of Chromosome 2, display particularly low levels of genetic differentiation between populations (Figure 3C), pointing to the potential for varying rates of recombination across the *P. modestum* genome. It is commonly recognized that recombination rate can be suppressed at different regions of the genome, a phenomenon typically characterized by fewer polymorphisms and non-random associations between alleles (Nachman, 2002; Renaut et al., 2013). In squamates, recombination rate has also been negatively correlated with distance to chromosome ends. (Schield et al., 2020). Interestingly, the *P. modestum* genomic regions displaying low levels of genetic divergence are similar to those regions displaying low levels of genetic divergence in *P. platyrhinos*, raising the question as to whether this property is preserved across taxonomic lineages in *Phrynosoma* (Farleigh et al., 2021). Further comparative examinations of *Phrynosoma* genome diversity are needed to understand broader patterns of genome evolution and their adaptive functions within the genus.

Gene ontology scans of known genes in windows of divergences identify multiple cases of functional interaction networks linked to a wide variety of cellular processes including translation initiation, mitosis, and promoting protein stability. In the South population, a functional interaction between DHX29, a translation initiation factor, and NOP14 is identified amongst potential genes in outlier windows under a divergence with gene flow model (Sweeney et al., 2021). Previous analyses of differentially expressed genes in *Anolis* lizards link genes involved in translation initiation with adaptation to climatic microhabitats (Akashi et al., 2016), which may be at play in the South population’s expansion into Northern Mexico. Additionally, a network of F-box proteins, including FBXL20 and FBXO11, along with MSH6, is found in shared divergence windows potentially under selection in allopatry (Mason & Laman, 2020); FBXO11 has been linked to thermal tolerance (Luna-Nevarez et al., 2021). No functional interactions are detected in genes linked to behavior, meaning current barriers to gene flow likely do not occur from a reproductive mismatch between the populations, as has been suggested for other regions of genetic divergence between *Anolis* species (Yang et al., 2020). This further supports the hypothesis that population divergence stems from allopatric forces rather than a sympatric mismatch.

To reduce the possibility of incorrectly identifying divergence windows due to non-selective genetic processes such as varying levels of recombination, the presence of directional selection within candidate windows is further tested through calculation of Tajima’s *D* (Figure 3B). Significantly negative values of Tajima’s *D* are detected in both divergence window scenarios in the North population when compared to the genomic background rate, indicating the presence of an excess of rare alleles that may have recently undergone a selective sweep. Tajima’s *D*, however, is also sensitive to non-selective demographic processes that act within a population to expose rare alleles, such as linkage, population expansion, and isolation-by-distance (Tajima, 1989). Our BPP analysis shows larger population sizes for both lineages compared to the ancestral population, and we cannot rule out the impact of recent population expansion on Tajima’s *D* along with heterozygosity metrics as mentioned above (Table S6).

While the population structure and demographic history of *P. modestum* characterized here present an important addition to the evolutionary history of phrynosomatid lizards, it is important to recognize the gap in sampling within our dataset as mentioned above. Some of the effects of this gap may be offset by the high coverage and number of loci recovered, in addition to the sampling distribution relative to the species range (Camargo et al., 2010, Leaché 2009). However, the addition of more individuals increases the accuracy of phylogenetic reconstruction more than the addition of more loci, and a broader resampling effort that includes regions omitted here may reveal further trends of migration or admixture between the two identified populations (Maddison & Knowles, 2006). Furthermore, although genetic sequencing of approximately 10x coverage yielded a large set of SNPs, the stringent filtering parameters applied reduced our dataset. Deeper coverage may be necessary for studies attempting to detect selective histories as performed here. Incorporation of morphological data to investigate correlations between genetic and phenotypic divergence will also further our understanding of *P. modestum* adaptation and potential speciation (McGreevey et al., 2024, Ogden & Thorpe, 2002). Despite these limitations, our investigation of genetic divergence and demography within *P. modestum* presents the species as one that can improve our understanding of the relative bounds of speciation under allopatric conditions.

## Data and code availability

Code and scripts to perform analyses are available on GitHub at https://github.com/kfarleigh/Modestum_WGS (to be made publicly available upon manuscript acceptance). Sequencing data are available as NCBI SRA BioProject XXXX (to be published upon manuscript acceptance).

## Author contributions

Designed study (CB, EC), performed laboratory and field work (CB, EC), performed statistical analyses (JA, KF, CB), participated in writing (JA, KF, EC, CB), project supervision and leadership (CB).

## Acknowledgments

We thank and acknowledge Dr. Jens Mueller for his assistance and for allowing us to use Miami University’s Redhawk high-performance computing cluster. We thank the following personnel and institutions for tissue loans: Jackson R. Roberts and William J. Stark (Sternberg Museum of Natural History), Lisa N. Barrow and J. Tomasz Giermakowski (Museum of Southwestern Biology), Carol L. Spencer, Jimmy A. McGuire and Rebecca Tarvin (Museum of Vertebrate Zoology), Travis LaDuc and David Cannatella (The University of Texas Austin Biodiversity Center), Greg Pandelis (The University of Texas Arlington), Carl S. Lieb, Eli Greenbaum, Jerry D. Johnson and Vicky Huang (The University of Texas El Paso), Adam D. Leaché and Peter Miller (Burke Museum of Natural History and Culture). We thank Tomas Flouri for assistance with BPP MSC-M analyses.

## Supplemental Materials

**Table S1.**
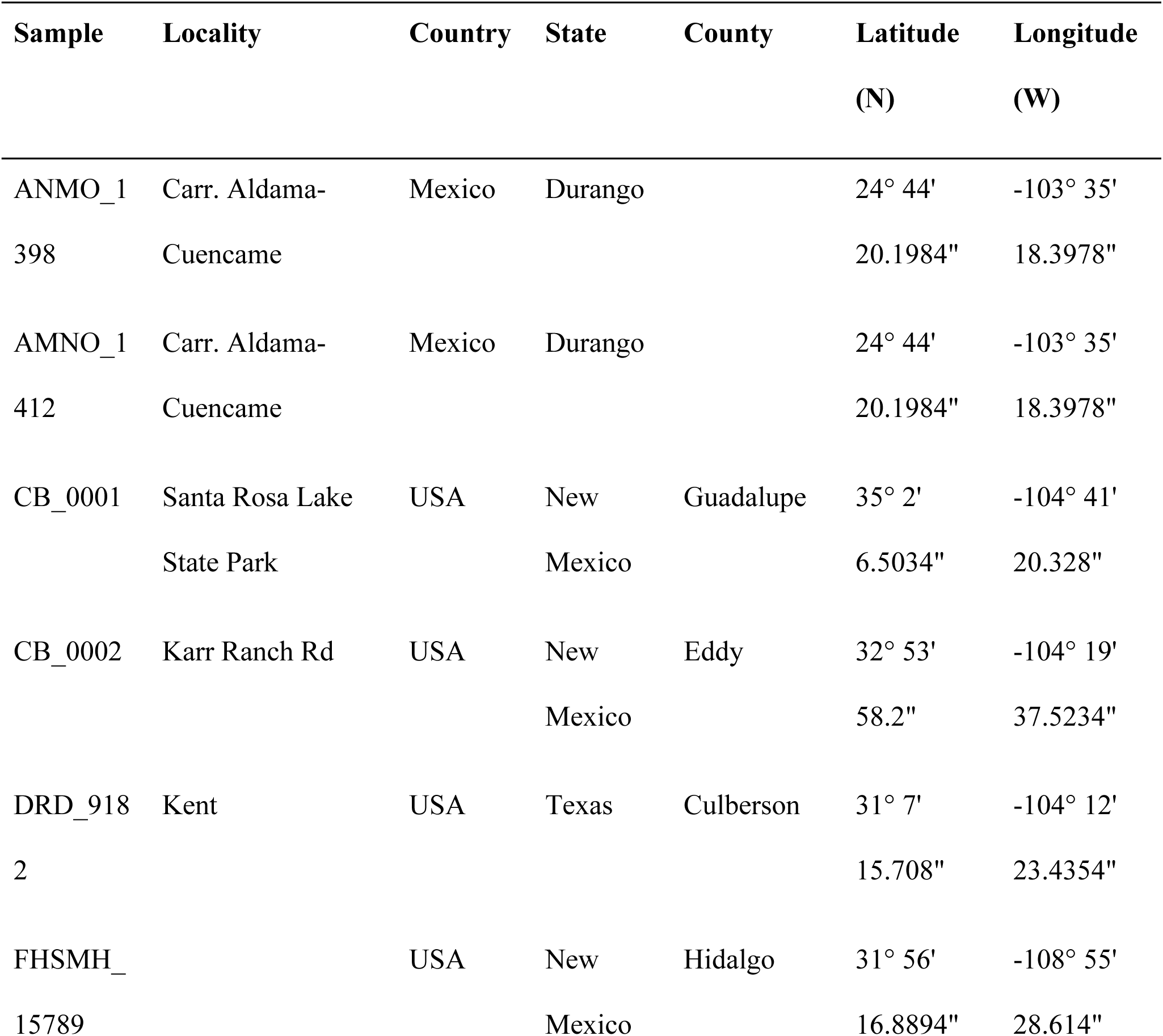
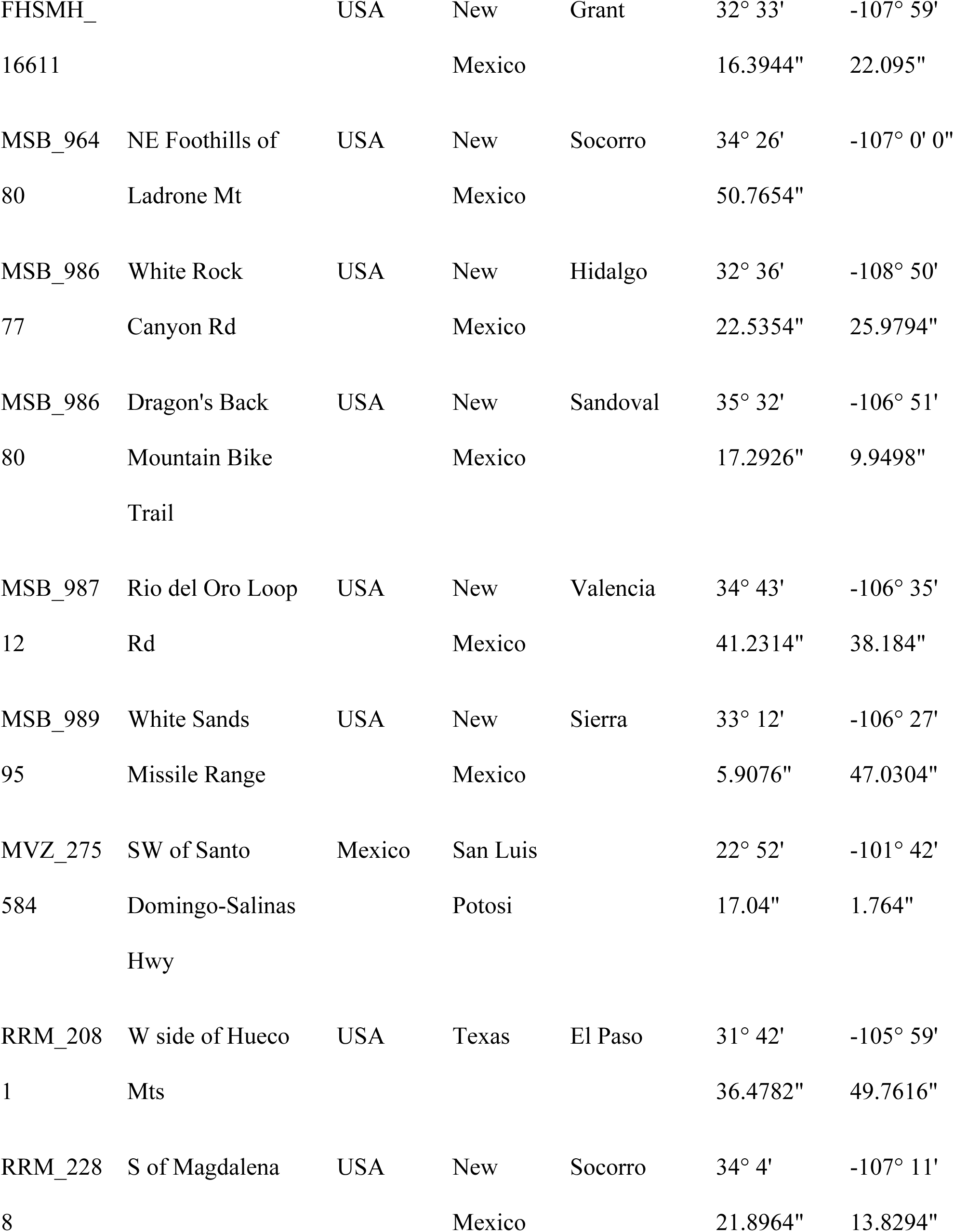
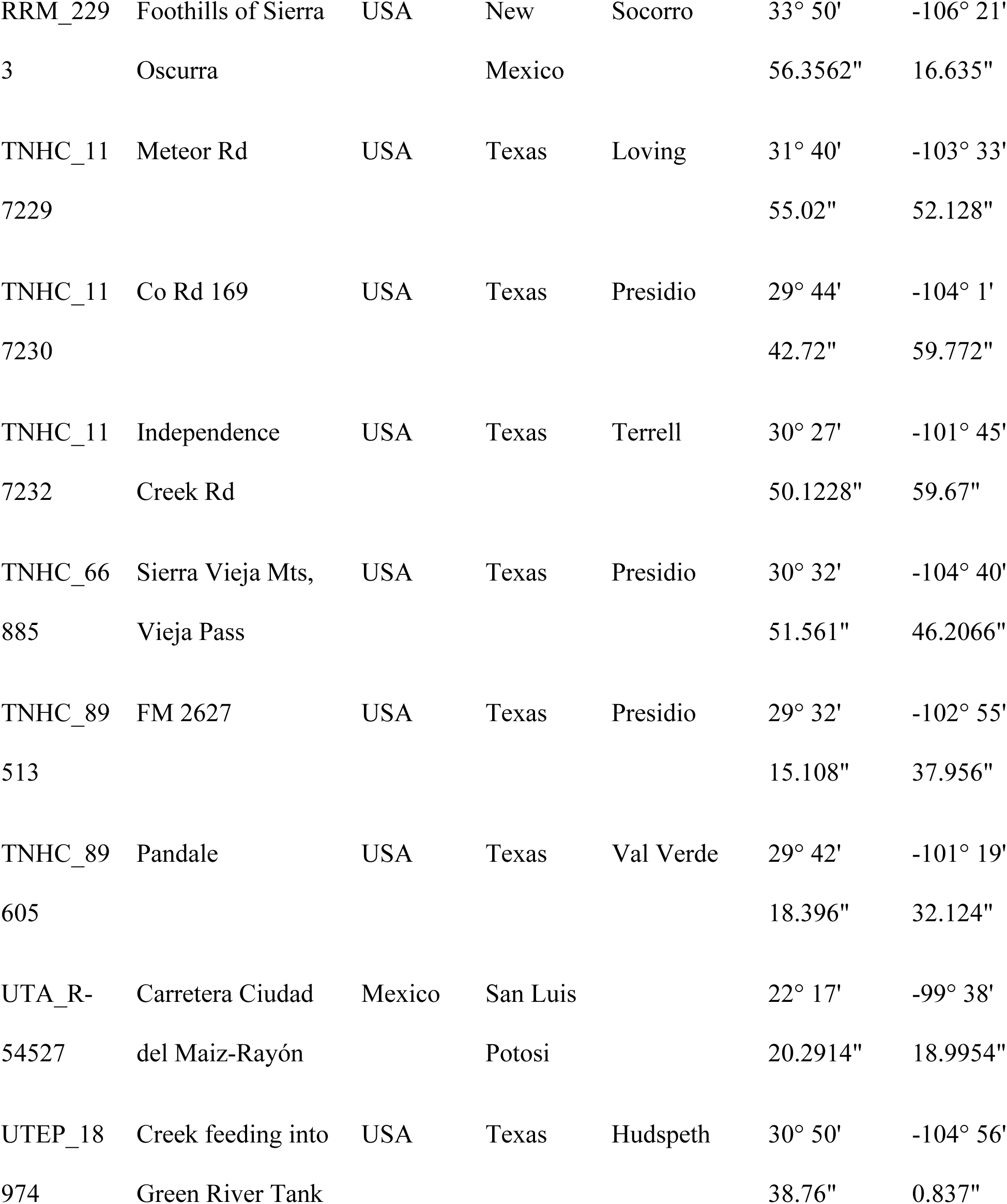
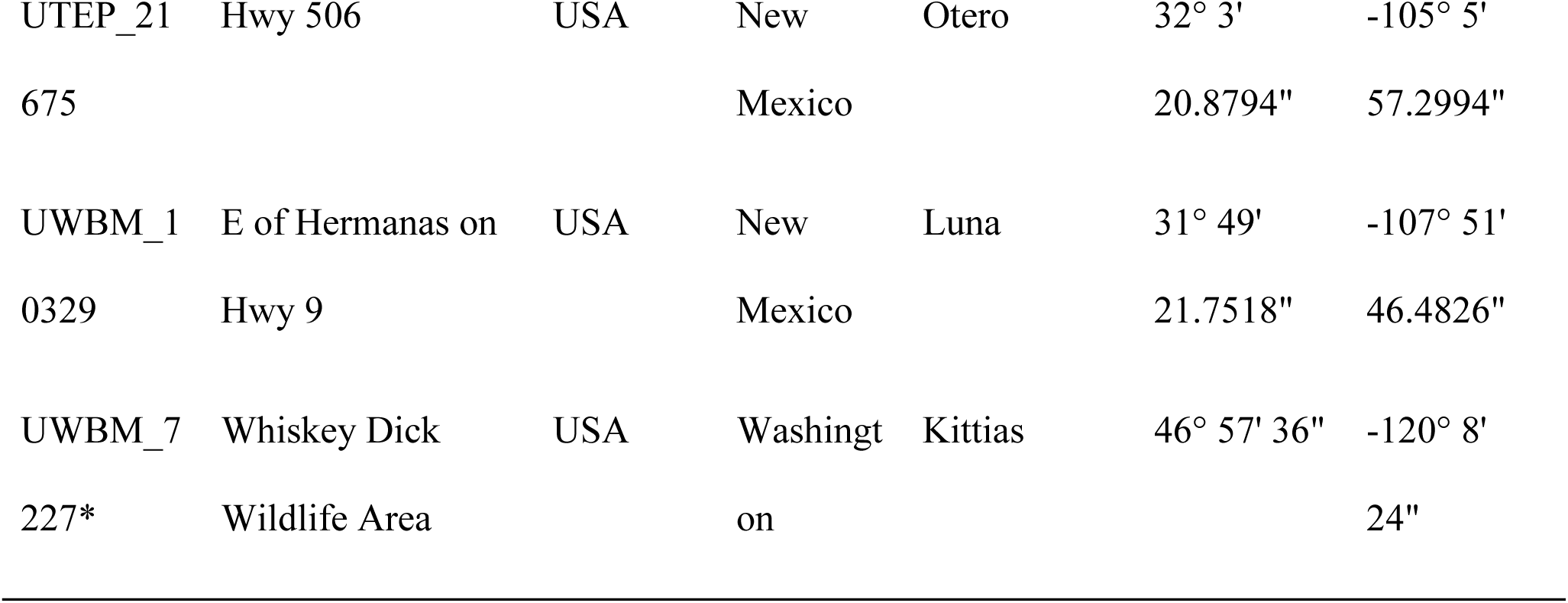
Description of sample areas for *Phrynosoma modestum* individuals, * indicates outgroup *Phrynosoma douglasii* individual.

**Table S2.** Description of sequencing statistics, species, and locality information for each sample included in this study.

| Sample | Total reads | Mapped reads | Mapping rate | Mean Depth |
| --- | --- | --- | --- | --- |
| ANMO_1398 | 156,840,714 | 147,530,115 | 94.06 | 10.49 |
| AMNO_1412 | 184,775,576 | 173,580,418 | 93.94 | 12.43 |
| CB_0001 | 170,056,600 | 160,173,985 | 94.19 | 11.25 |
| CB_0002 | 1561,685,84 | 147,168,023 | 94.24 | 10.3 |
| DRD_9182 | 216,434,726 | 203,181,419 | 93.88 | 14.27 |
| FHSMH_15789 | 124,752,768 | 116,221,604 | 93.16 | 8.39 |
| FHSMH_16611 | 136,770,378 | 128,745,895 | 94.13 | 9.23 |
| MSB_96480 | 165,605,308 | 155,094,579 | 93.65 | 11.07 |
| MSB_98677 | 167,583,434 | 157,633,034 | 94.06 | 11.08 |
| MSB_98680 | 144,175,988 | 135,477,985 | 93.97 | 9.81 |
| MSB_98712 | 154,081,250 | 144,378,927 | 93.7 | 10.87 |
| MSB_98995 | 146,676,728 | 137,507,279 | 93.75 | 10.08 |
| MVZ_275584 | 168,625,166 | 156,910,490 | 93.05 | 10.95 |
| RRM_2081 | 116,843,970 | 31,610,765 | 27.05 | 9.97 |
| RRM_2288 | 139,379,022 | 126,885,046 | 91.04 | 7.73 |
| RRM_2293 | 101,982,616 | 94,767,735 | 92.93 | 7.26 |
| TNHC_117229 | 615,42,144 | 57,473,958 | 93.39 | 4.66 |
| TNHC_117230 | 162,635,040 | 152,713,095 | 93.9 | 11.24 |
| TNHC_117232 | 146,416,076 | 137,718,557 | 94.06 | 10.21 |
| TNHC_66885 | 165,982,074 | 155,581,482 | 93.78 | 11.19 |
| TNHC_89513 | 166,913,824 | 156,592,293 | 93.82 | 11.71 |
| TNHC_89605 | 141,893,238 | 133,213,806 | 93.88 | 9.79 |
| UTA_R-54527 | 171,487,628 | 161,438,404 | 94.14 | 11.61 |
| UTEP_18974 | 105,512,680 | 98,440,779 | 93.3 | 7.46 |
| UTEP_21675 | 97,981,490 | 90,934,861 | 92.81 | 6.82 |
| UWBM_10329 | 149,387,068 | 140,549,550 | 94.08 | 9.86 |
| UWBM_7227 | 172,797,520 | 158,880,949 | 91.95 | 11.4 |

**Table S3.** Captus assembly statistics for each sample included in this study.

| Sample | Assembled<br>contigs | Mean Depth | Extracted loci | % genes<br>recovered |
| --- | --- | --- | --- | --- |
| ANMO_1398 | 1,189,176 | 7.70 | 5,090 | 93.0 |
| AMNO_1412 | 1,166,892 | 8.59 | 5,092 | 93.1 |
| CB_0001 | 809,073 | 9.83 | 5,093 | 93.1 |
| CB_0002 | 861,544 | 9.27 | 5,090 | 93.0 |
| DRD_9182 | 1,027,038 | 9.99 | 5,099 | 93.2 |
| FHSMH_15789 | 1,253,903 | 8.27 | 5,063 | 92.5 |
| FHSMH_16611 | 905,106 | 8.64 | 5,092 | 93.1 |
| MSB_96480 | 951,329 | 9.25 | 5,032 | 92.0 |
| MSB_98677 | 809,992 | 10.00 | 5,092 | 93.1 |
| MSB_98680 | 831,588 | 8.96 | 5,088 | 93.0 |
| MSB_98712 | 819,818 | 9.84 | 5,092 | 93.1 |
| MSB_98995 | 815,463 | 8.69 | 5,090 | 93.0 |
| MVZ_275584 | 1,119,828 | 8.70 | 5,097 | 93.1 |
| RRM_2288 | 1,399,739 | 8.59 | 4,937 | 90.2 |
| RRM_2293 | 1,556,661 | 7.47 | 4,901 | 89.6 |
| TNHC_117229 | 1,945,444 | 5.27 | 4,501 | 82.3 |
| TNHC_117230 | 1,023,242 | 8.43 | 5,087 | 93.0 |
| TNHC_117232 | 977,575 | 8.29 | 5,088 | 93.0 |
| TNHC_66885 | 999,003 | 8.65 | 5,091 | 93.1 |
| TNHC_89513 | 994,478 | 9.09 | 5,101 | 93.2 |
| TNHC_89605 | 1,028,500 | 7.98 | 5,091 | 93.0 |
| UTA_R-54527 | 819,571 | 9.13 | 5,101 | 93.2 |
| UTEP_18974 | 1,491,906 | 7.05 | 4,928 | 90.1 |
| UTEP_21675 | 1,820,505 | 7.07 | 4,873 | 89.1 |
| UWBM_10329 | 2,039,949 | 6.23 | 4,020 | 73.5 |
| UWBM_7227 | 2,091,819 | 6.21 | 4,811 | 87.9 |

**Table S4.** Parameter estimates from BPP MSC-M analyses. Results are from four independent runs. Two migration events were specified in the model. Values of theta and tau are multiplied by 1000. A = North, B = South.

| Parameter | Mean | 95% HPD lower | 95% HPD upper |
| --- | --- | --- | --- |
| $\theta_A$ | 12.3 | 12.2 | 12.4 |
| $\theta_B$ | 10.6 | 10.3 | 10.8 |
| $\theta_{AB}$ | 3.2784 | 3.184 | 3.368 |
| $\theta_{\text{root}}$ | 19.6 | 19 | 20.2 |
| $\tau_{\text{root}}$ | 5.9882 | 5.902 | 6.072 |
| $\tau_{AB}$ | 3.1152 | 3.085 | 3.145 |
| $\omega_{A \rightarrow B}$ | 1.416 | 0.7804 | 2.0779 |
| $\omega_{B \rightarrow A}$ | 0.1263 | 0.0156 | 0.2483 |

**Table S5.**
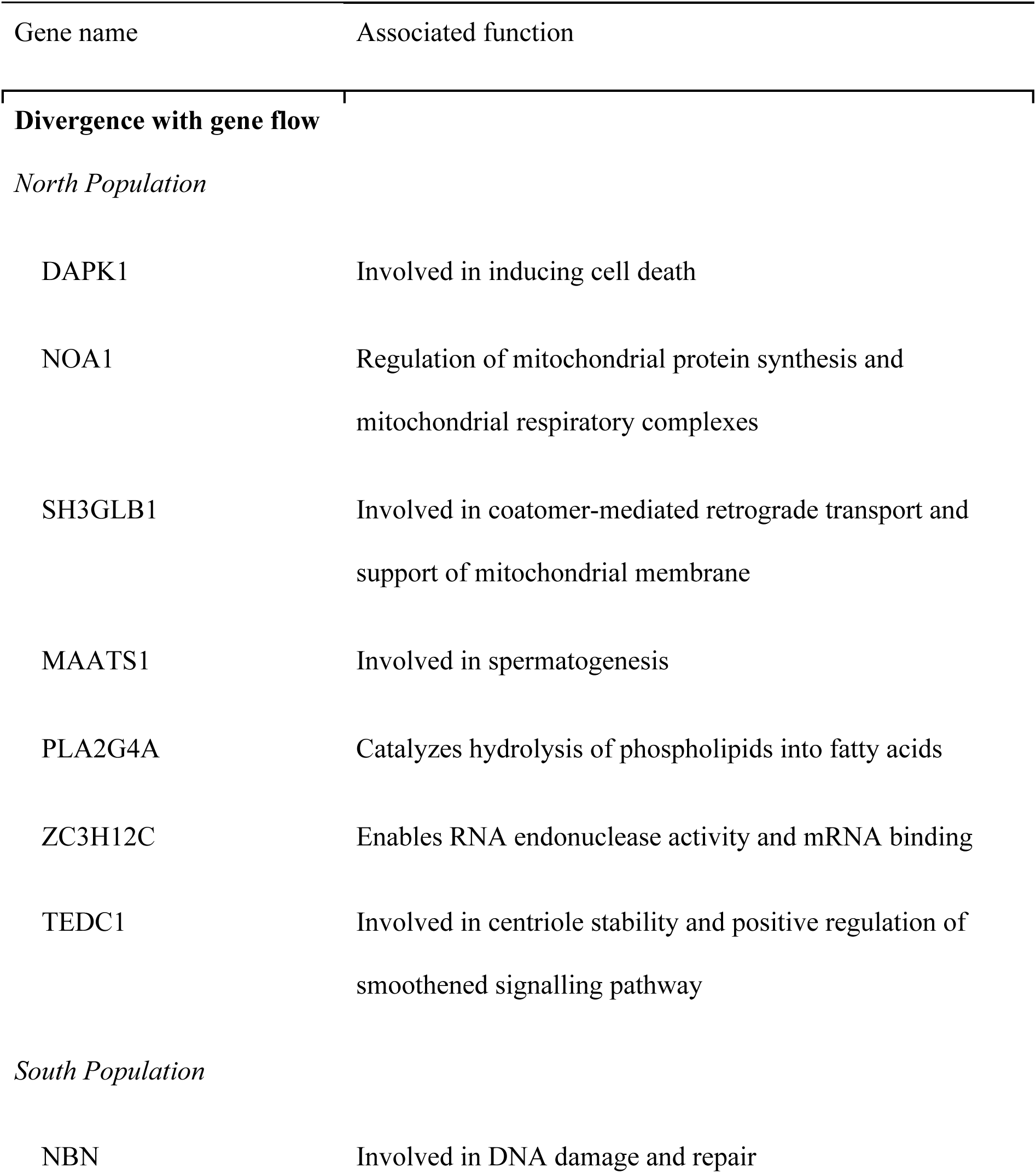
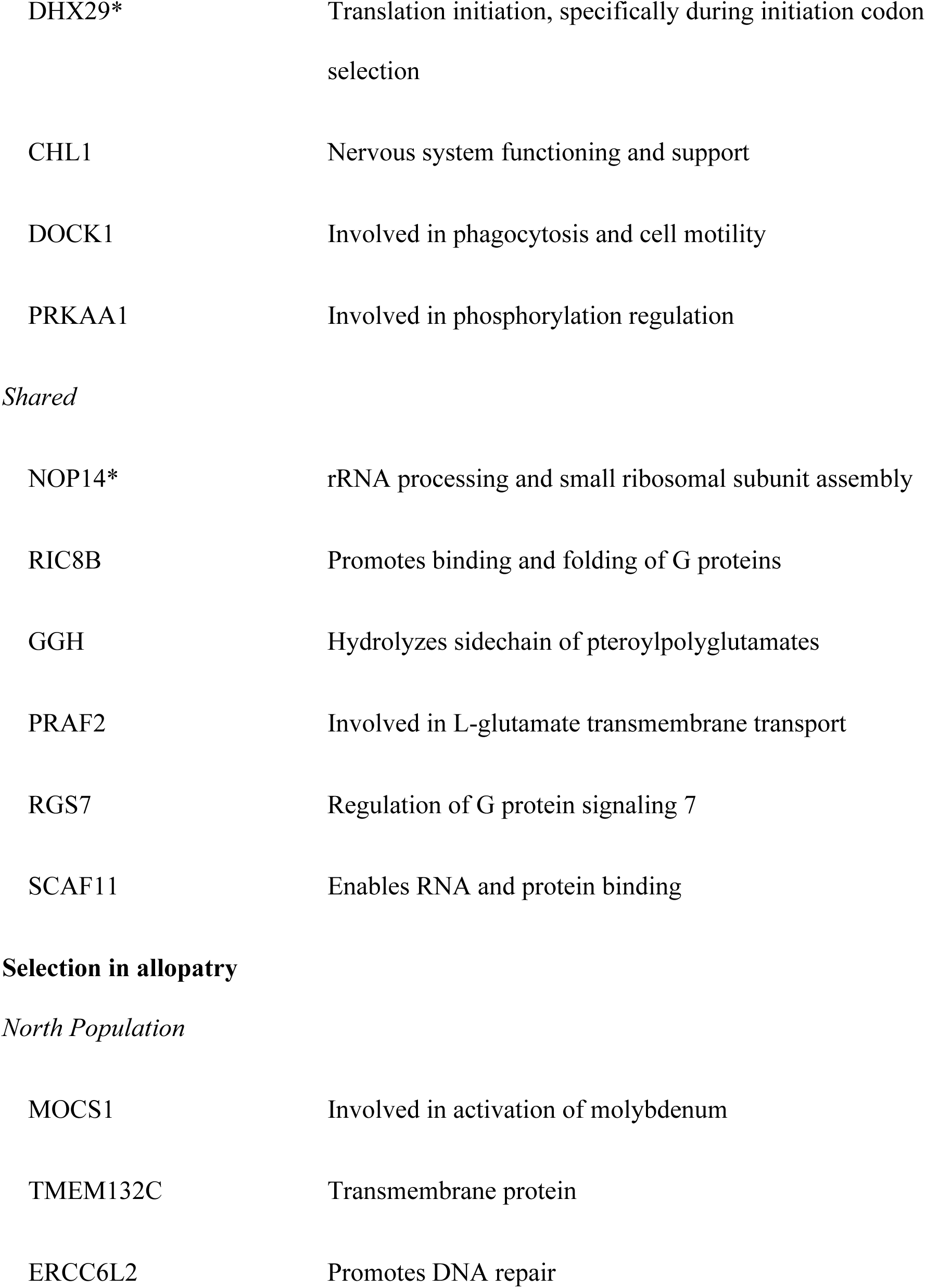
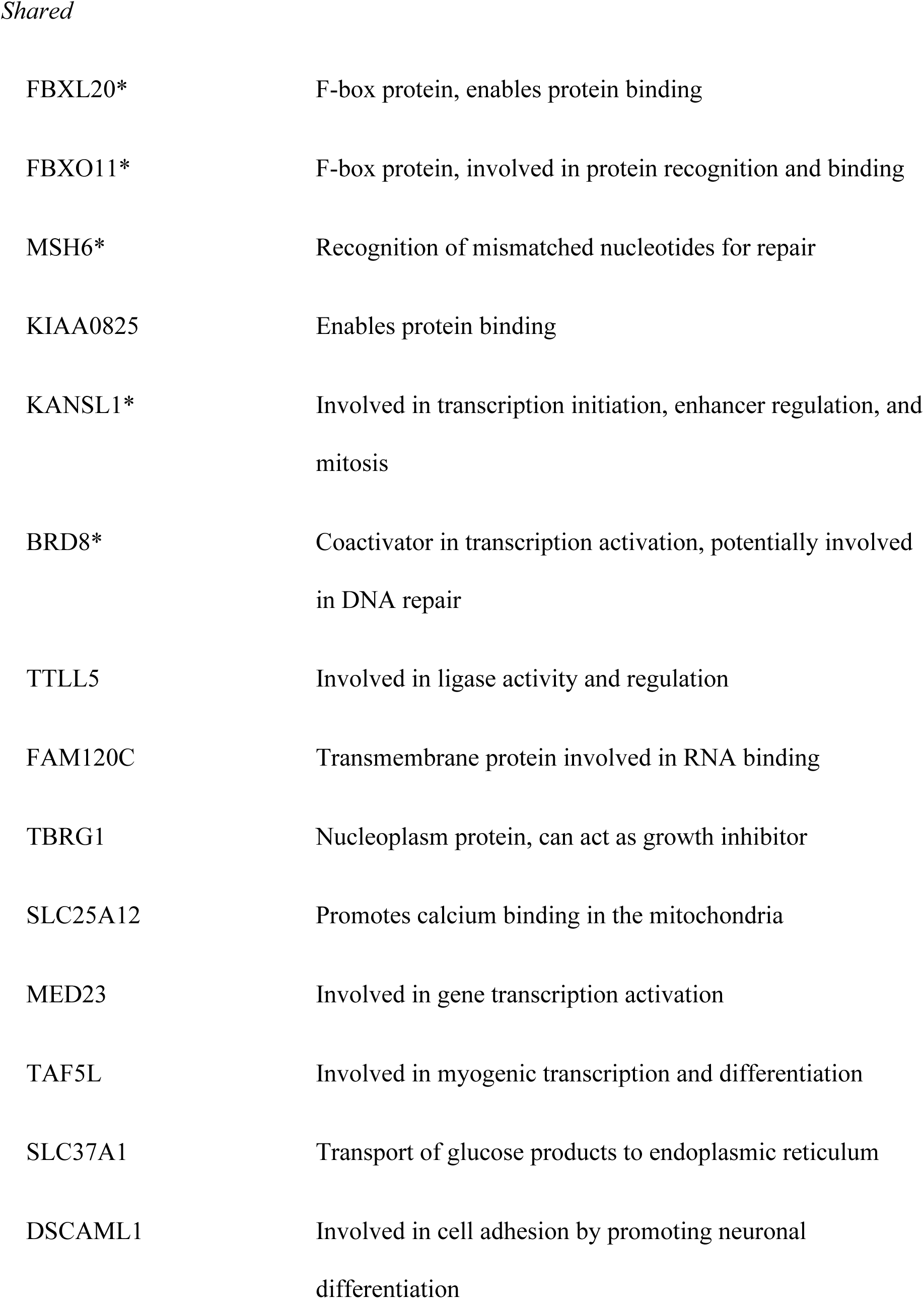
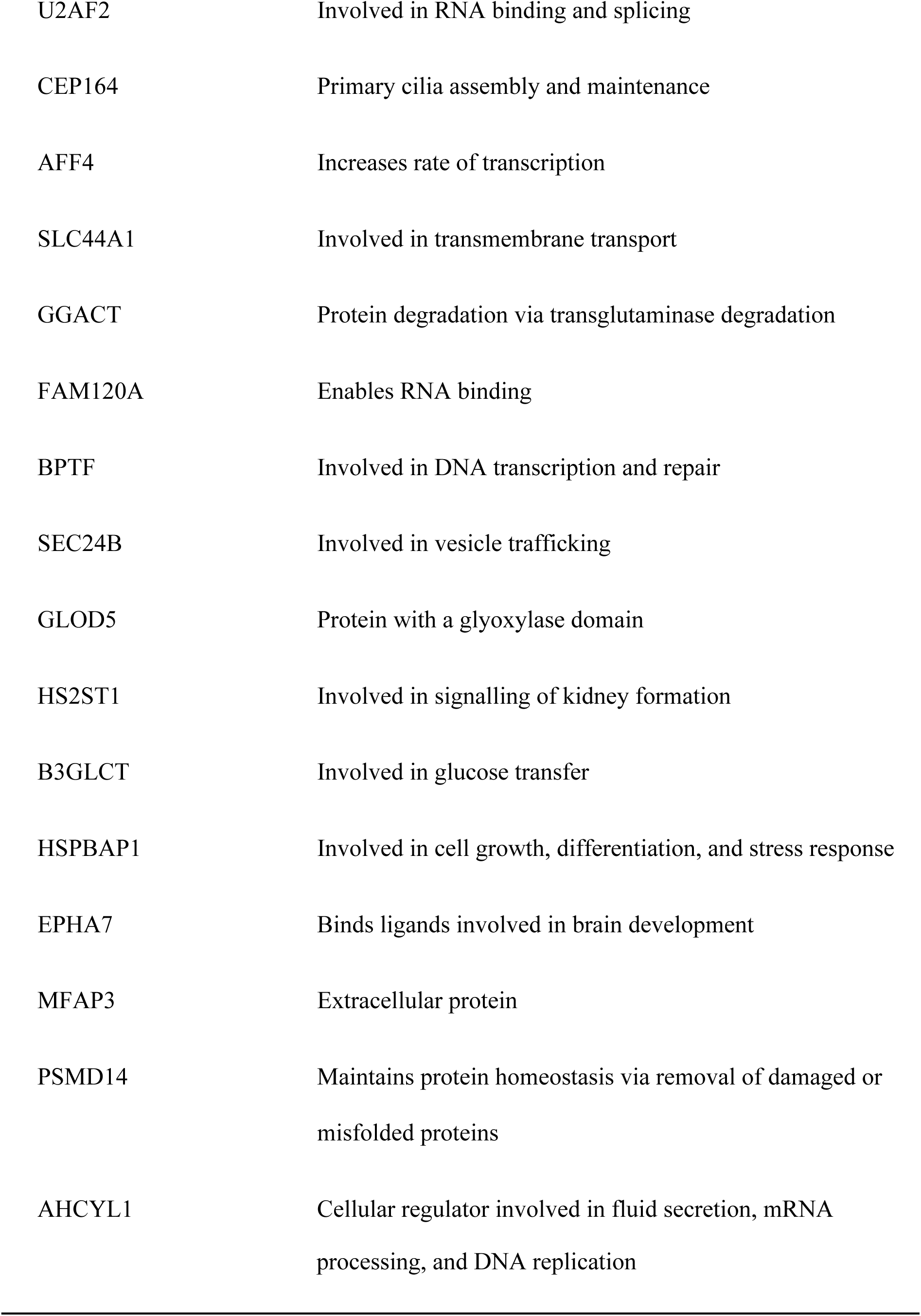
Molecular functions of genes identified within 10kb windows of divergence. * indicates a gene’s presence in a functional interaction network.

**Table S6.** The proportion of heterozygous loci for each individual and their population assignment.

| Sample | Proportion of heterozygous loci | Population |
| --- | --- | --- |
| FHSMH_16611 | 0.246448 | North |
| MSB_98680 | 0.233379 | North |
| MSB_96480 | 0.245329 | North |
| TNHC_66885 | 0.329061 | North |
| MSB_98677 | 0.246916 | North |
| TNHC_89605 | 0.345256 | North |
| DRD_9182 | 0.32303 | North |
| UWBM_10329 | 0.257765 | North |
| CB_0002 | 0.278595 | North |
| CB_0001 | 0.268774 | North |
| RRM_2288 | 0.233891 | North |
| MSB_98995 | 0.237231 | North |
| MSB_98712 | 0.258095 | North |
| FHSMH_15789 | 0.271081 | North |
| UTEP_21675 | 0.284863 | North |
| UTEP_18974 | 0.306929 | North |
| TNHC_89513 | 0.365027 | North |
| TNHC_117229 | 0.308165 | North |
| RRM_2293 | 0.263945 | North |
| TNHC_117232 | 0.346356 | North |
| TNHC_117230 | 0.358369 | North |
| ANMO_1398 | 0.447393 | South |
| AMNO_1412 | 0.454903 | South |
| MVZ_275584 | 0.443618 | South |
| UTA_R-54527 | 0.343364 | South |

**Figure S1.**
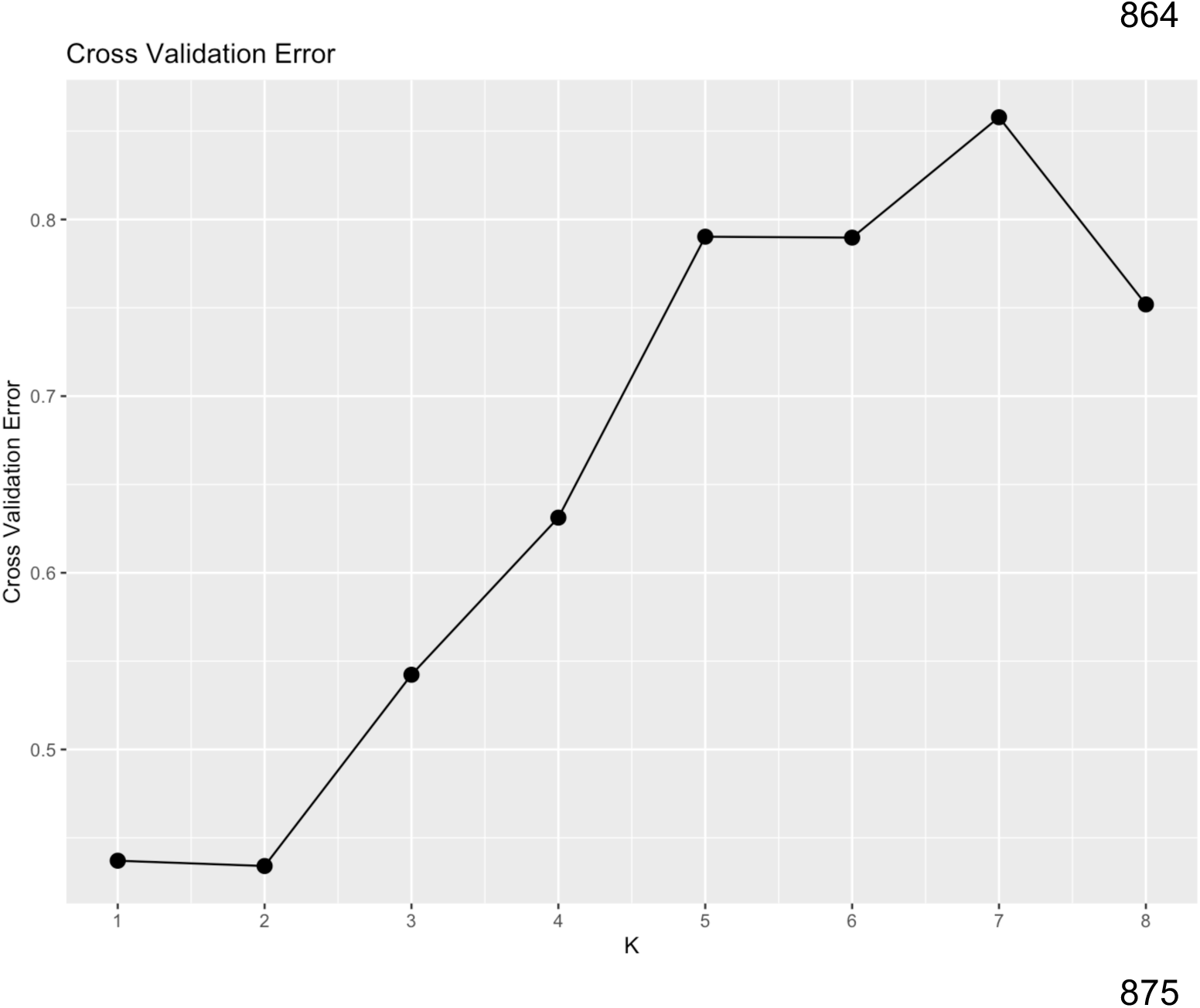
Cross-validation error for ADMIXTURE analyses for values of K = 1 - 8. K = 2 minimizes cross-validation error at 0.434.

**Figure S2.**
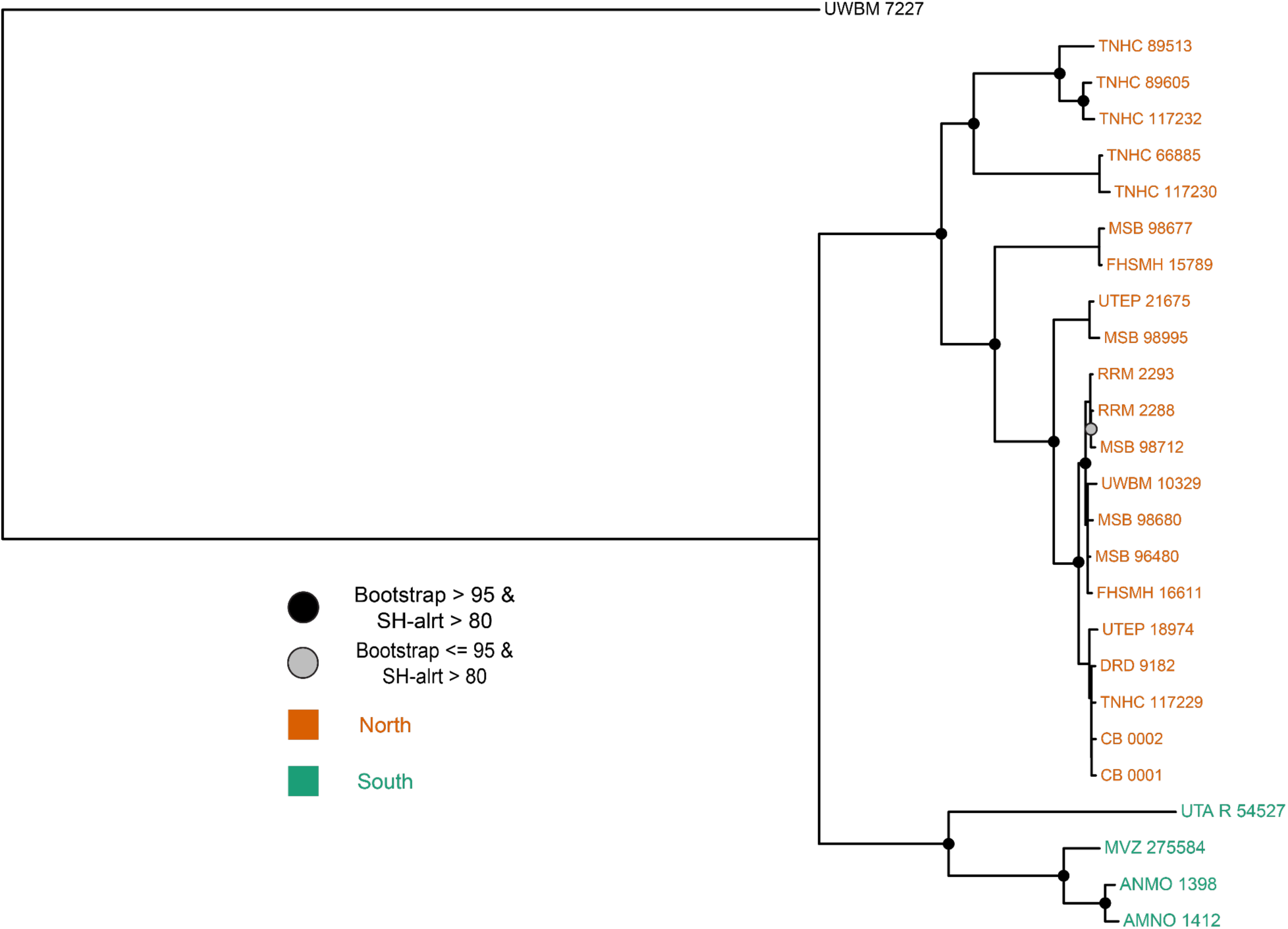
Maximum-likelihood (ML) genealogy of whole mitochondrial DNA (mtDNA) genome sequences using IQ-TREE. Mitochondrial genomes were extracted using captus, with the *Phrynosoma blainvilli* mtDNA genome as the reference. Results are generally concordant with the tetrapod ultra-conserved element genealogy presented in the main text.

